# Long-term two-photon imaging of spinal cord in freely behaving mice

**DOI:** 10.1101/2022.01.09.475306

**Authors:** Furong Ju, Wenling Jian, Yaning Han, Tianwen Huang, Jin Ke, Zhiheng Liu, Shengyuan Cai, Nan Liu, Liping Wang, Pengfei Wei

**Affiliations:** Shenzhen Key Lab of Neuropsychiatric Modulation and Collaborative Innovation Center for Brain Science, Guangdong Provincial Key Laboratory of Brain Connectome and Behavior, CAS Center for Excellence in Brain Science and Intelligence Technology, Brain Cognition and Brain Disease Institute (BCBDI), Shenzhen-Hong Kong Institute of Brain Science-Shenzhen Fundamental Research Institutions, Shenzhen Institute of Advanced Technology, Chinese Academy of Sciences, Shenzhen, China; University of Chinese Academy of Sciences, Beijing, China; Department of Anesthesiology, Shenzhen Second People’s Hospital, The First Affiliated Hospital of Shenzhen University, Shenzhen, China

**Author notes:** Corresponding author: Pengfei Wei and Liping Wang. Contributed equally (Furong Ju, Wenling Jian, Yaning Han).

**Keywords:** spinal cord, dorsal horn, two-photon microscopy, neuropathic pain, natural behavior, sensory perception, in vivo, cutaneous stimulation, subcellular resolution, protective motor behavior

## Abstract

The spinal cord is critical to the perception of peripheral information under sensory-guided motor behaviors in health and disease. However, the cellular activity underlie spinal cord function in freely behaving animals is not clear. Here, we developed a new method for imaging the spinal cord at cellular and subcellular resolution over weeks under naturalistic conditions. The method involves an improved surgery to reduce spinal movement, and the installation of a miniaturized two-photon microscope to obtain high-resolution imaging in moving mice. In vivo calcium imaging demonstrated that dorsal horn neurons show a sensorimotor program-dependent synchronization and heterogeneity under distinct cutaneous stimuli in behaving mice. The long-term imaging of sensory neurons revealed that in the spinal cord, healthy mice demonstrated stereotyped responses. However, in a neuropathic pain model, plasticity changes and neuronal sensitization were observed. We provide a practical method to study the function of spinal cord on sensory perception and disorders in freely behaving mice.

## Main

Afferent spinal neurons are fundamentally responsible for the integration and transmission of messages from peripheral receptors to the brain ^1–4^. Recent research assessing mechanoreceptors and thermoceptors in sensory neurons has revealed that the spinal cord plays a key role in coding of perceptual information and multimodal responses ^5–8^. Traditionally, the role of different neurons within circuits encoding and relaying somatic sensations has largely been determined according to their distinct morphological and electrophysiological properties ^4,9–12^. While these studies have generated critical knowledge advancing the field, studies have remained restricted to anesthetized animals. Anesthesia can interfere with animal recognition, perception, and motor control, and recordings made under anesthesia fails to accurately explain how the somatosensory system encodes a wide range of tactile stimuli^13–15^, and the critical aspects of spinal cord sensorimotor processing remain poorly understood. In addition, spinal cord circuit tracing studies have revealed important roles for sensory-guided motor behaviors, such as reflex-defenses and coping responses induced by noxious stimuli^16^, but these genetic, electrophysiological and pharmacologic approaches still insufficiently describe the spatial and temporal neuronal functional patterns that underlie external stimuli-guided different motor behaviors. The recent real-time imaging methods for interrogation of neuronal activity provide a new insight into how heterogeneous cell types encode noxious and innocuous stimulations from peripheral inputs^5,8^. However, elucidating the morphological and functional properties of spinal cord neurons continues to be a major challenge in free-behaving mice due to the lack of suitable optical technology.

Optical technology combined with behavioral research provides a comprehensive sampling method. This can provide the capacity for high spatial and temporal resolution during long-term observation windows, as well as multidimensional behavioral tracking of animal behavior ^17,18^. In recent years, the incorporation of calcium imaging into behavioral research has deepened our knowledge about how cerebral neurons encode, store, and modify incoming and outgoing information in awake and unrestrained animals ^17,18^. Current miniaturized one-photon microscopy techniques allow for high-speed fluorescence calcium imaging and a large field-of-view (FOV) of the spinal cord, yet retain a number of limitations. First, the superior contrast and spatial resolution restricts the performance of miniaturized one-photon microscopy. Second, high neuron density and cell infiltration may result in the mixing of signals or loss of focus ^19^. In addition, motion artifacts during free movement can cause loss of image authenticity certification ^5,8,14,20^. In an effort to provide a fluorescence imaging approach based on miniaturized two-photon microscopy, we developed a spinal fusion method in mice, which minimizes spinal movement and allows long-term imaging of spinal cord neurons at cellular and subcellular resolution in free-behaving states. Using this spinal fusion alongside a removable miniaturized two-photon imaging piece, we provide continuous optical access to the mouse spinal. This method facilitates in vivo studies of dorsal horn-related disorders and neuropathies at both the cellular and subcellular levels.

## Results

### Two-photon imaging in spinal cord of freely behaving mice

To image the spinal cord in freely moving mice, we designed a custom implantable spinal chamber **(Fig. 1 A)** and a surgical procedure to enable in vivo imaging of miniaturized two-photon microscopy **(Fig. 1 B–C)**. The spinal chamber was composed of two stainless steel grooves that were implanted under the transverse process of vertebrae that could be firmly attached to the spinal column and minimized motion artifacts **(Supplementary fig. 1 A–B)**. The surgical procedure involved removal of the muscle and tendon tissue from the dorsal laminae over three vertebrae (L3/4/5 or L4/5/6), then fusing these vertebrae by clamping them on either side with a stainless steel groove held by stereotaxic apparatus **(Supplementary fig. 1 A–B)**. Finally, a dorsal laminectomy was performed using a high-speed dental drill. We maintained the clamping pressure by placing a 4–5 mm glass coverslip on top (which further minimized motion artifacts), applying Kwik-Sil silicone elastomer over the spinal cord, sealing the chamber by filling the gap with a combination of cyanoacrylate and glue, and finally attaching a miniaturized two-photon microscope to the animal’s back through a headpiece which could be removed and replaced **(Fig. 1 C, Supplementary fig. 1 C–G)**. The entire assembled piece was small and light enough to be borne by the mice **(Fig. 1 D–E)**. Before surgical preparation, and 30 days afterwards, we quantified the animals’ locomotor performance in a 5-min open field test (OFT). Neither mice with or without miniature two-photon microscopy (mTPM) showed a significant difference in total moving distance compared to sham mice (mice before surgery), and both groups showed a significant decrease in average speed after compared to sham mice **(Fig. 1 F–G)**.

**Figure 1.**
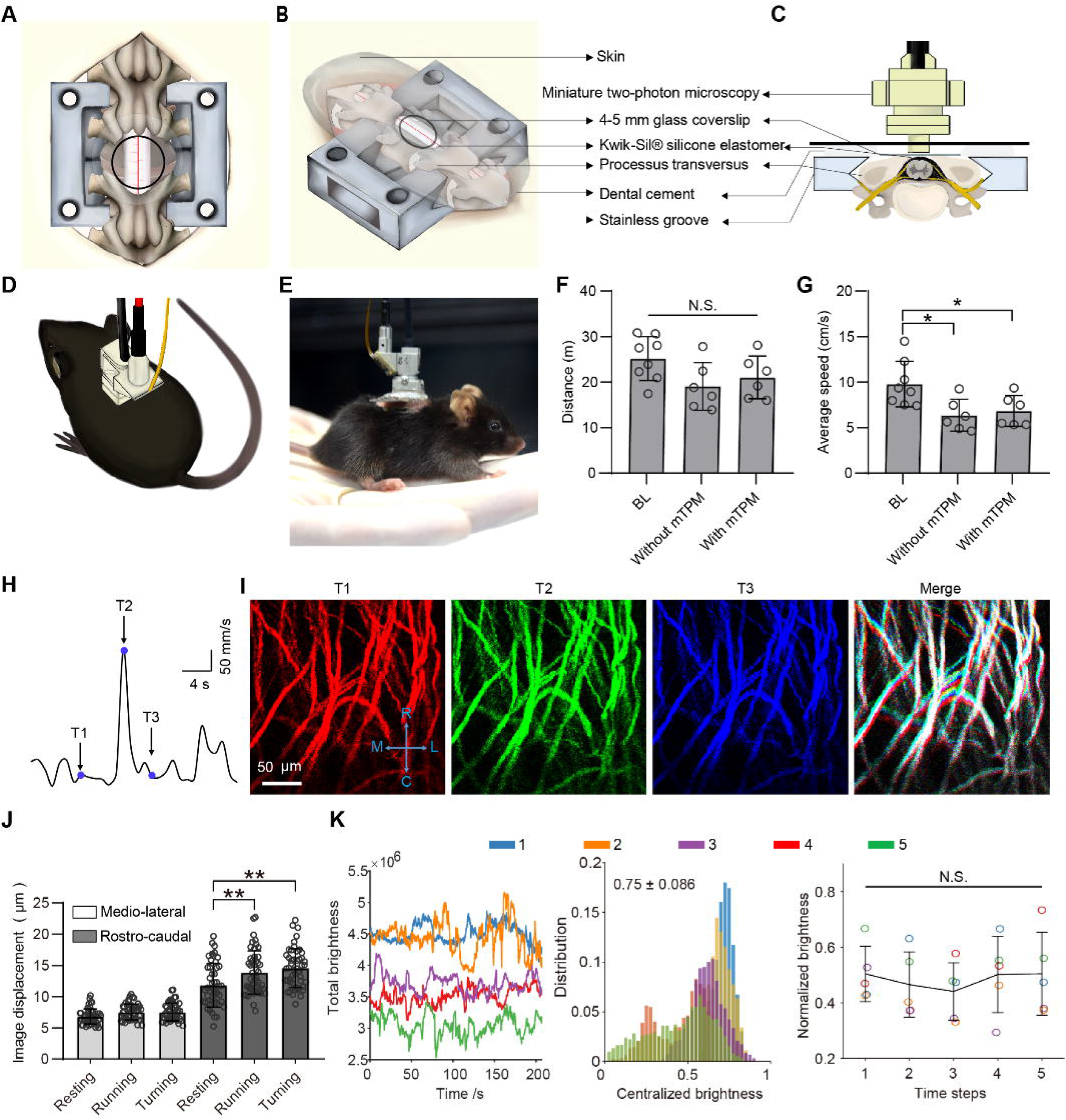
Performance of FHIRM-TPM compared with an imaging chamber for optical access to the spinal cord in freely behaving mice. **A)** Image of custom-designed implantable spinal chamber placed under the transverse process at lumbar level L3/4/5 or L4/5/6. **B–C)** Schematics showing the implanted imaging chamber and the two-photon microscope objective in cross-section. **D–E)** Illustration and photograph representing a mouse wearing the FHIRM-TPM headpiece. **F–G)** Total travel distance (left) and average speed (right) in the locomotor test for mice before surgery at baseline (BL), with chamber only (without mTPM), or mounted with mTPM (N=6–8 mice per group; *N.S. P*>0.05, \**P*<0.05). **H–I)** Mouse movement speed (black trace) during a 30 s recording period (left) and representative images of spinal cord axons at the initial frame (T1), the frame corresponding to the maximum speed of the mouse (T2), and the frame after running bouts (T3) (right). Image displacement occurred mainly in the rostrocaudal direction (Merge). Scale bars, 50 mm/s and 4 s (top); 50 mm (bottom). **J)** Population data showing image displacement in mediolateral or rostrocaudal directions occurred when the mouse was resting, running, and turning (N=6 mice per group; \*\**P* < 0.01). **K)** Total brightness of all frames traced for 200 s from 5 mice showing image displacements in the dorsoventral direction. Left: brightness changes of axons in real-time imaging of 5 typical mice; middle: MS-SSIM index showing the distribution of images of all frames within 200 s from 5 mice; right: normalized brightness is depicted as a probability distribution of image displacements in the dorsoventral direction (N=5 mice; each curve represents data from an individual animal). Note that there is almost no image loss in the dorsoventral direction. *N.S. P* > 0.05. FHIRM-TPM, fast high-resolution miniature two-photon microscopy; mTPM, miniaturized two-photon microscopy; MS-SSIM, multiscale structural similarity.

To quantify the motion artifacts caused by physical activity or free movement of the awake mice, we implanted spinal cord windows and FHIRM-TPM in transgenic Thy1-GFP mice with sparsely labelled axons. In free-moving mice, there were image displacements due to different moving speeds **(Fig. 1 H–I)** and increased in amplitude during steady running or turning. These remained largely confined to the rostrocaudal direction. Meanwhile, displacements in the mediolateral direction were small, and there were no significant differences in resting, turning, or running **(Fig. 1 J**; image displacement in the mediolateral direction: Resting 6.8±1.2 μm, Running 7.5±1.3 μm, Turning 7.6±1.4 μm; image displacement in the rostrocaudal direction: Resting 11.8±3.5 μm, Running 13.9±3.5 μm, Turning 14.6±3.1 μm; data shown as mean ± standard error of the mean, SEM). Next, we quantified displacement in the dorsoventral direction during imaging, which we determined to be more critical as this could cause the overall image to be lost. We analyzed the brightness distribution and consistency of axons in real-time imaging. The brightness of axons was not lost and showed no changes during the continuous imaging process and the multiscale structural similarities (MS-SSIMs) showed that frames from one mouse maintained a high degree of consistency and similarity, showing the average MS-SSIM of 0.75±0.086 **(Fig. 1 K)**. These results show that the structure of axons in the spinal cord of free-moving mice can be reliably imaged over extended periods through an implanted vertebral window and a miniaturized two-photon microscope, without compromising motor functions.

### Calcium imaging of spinal dorsal horn neurons in free-moving mice

In contrast to the well-characterized anatomy and function of the primary sensory neurons in the DRG, our understanding of sensory neurons in the spinal cord remains limited ^15^. To address this, we next performed in vivo Ca^2+^ imaging to examine the somatic activity of sensory neurons expressing the Ca^2+^ indicator GCaMP6s in the spinal cord of free-moving mice using a miniaturized two-photon microscope. To minimize the potential effects of surgery-related inflammation induced by spinal window implantation, the first imaging was performed 30 days after surgery. With a FOV of 420×420 μm^2^, we resolved 54.2±5.3 neurons in a single focal plane at a frame rate of 4.84 Hz at 512×512 pixels. In the quiet-awake-fixed mice, spontaneous Ca^2+^ transients of dorsal horn neurons (349 neurons from 7 mice, 51 spontaneous active neurons) were clearly imaged with a few active populations and low response amplitudes **(Fig. 2 A)**. In contrast, marked spontaneous Ca^2+^ transients were observed compared to the pre-awake-fixed state in free-moving mice (97 spontaneous active neurons, **Supplementary Movie 1**), and significant increases in the active neuron population and maximum response amplitudes (*ΔF*/*F0*) were found across all imaging sessions (active population: 11.3±6.4% for Awake versus Free 26.6±6.1%, *ΔF/F0*: 1.7±0.1 for Awake versus Free 3.2±0.2; data shown as mean ± SEM). To further assess the spontaneous Ca^2+^ transients of dorsal horn neurons in free-moving mice, the intensity of the cellular calcium signal of activated neurons was traced and labelled **(Fig. 2 D–E)**. We evaluated the synchronous calcium release pattern for a 10-min open field test in free-moving mice. Results of this indicated that spontaneous Ca^2+^ transients have a high degree of synchronization in free-moving mice, which differs from the spontaneous activity of neurons in the brain **(Fig. 2 F–G)**. To determine whether these spontaneous Ca^2+^ transients of dorsal horn neurons were related to motion or sensation, we further explored the relationship between mouse behavior and the corresponding neuron calcium intensity. As shown at the bottom of **Fig. 2 H**, we recorded the calcium activity of 59 neurons from one mouse under free-moving conditions and when with air-puff stimuli. First, we employed principal component analysis (PCA) to extract the principal linear combinations of high-dimensional neural activities, which we hypothesized could be correlated with the speed of the mouse. Nineteen PCs were selected and sorted based on the explanation of 90% variance of neural activities (**Fig. 2 H**, middle). The locomotion speed of the mouse and start time of air stimuli are shown at the top of **Fig. 2 H**. The correlation coefficients (CCs) of five mice were mostly distributed in the range of −0.3 to 0.3, which verified that correlations between neural activity and speed were low **(Fig. 2 I**). We next partitioned the speed into three levels to investigate the activity of neurons at different speed levels with air-puff stimuli. We found that the activity of neurons increased during and after airpuff compared to free and before air-puff (**Fig. 2 J**). Furthermore, neuronal activity was significantly increased during air puff, regardless of the speed levels **(Fig. 2 K**).

**Figure 2.**
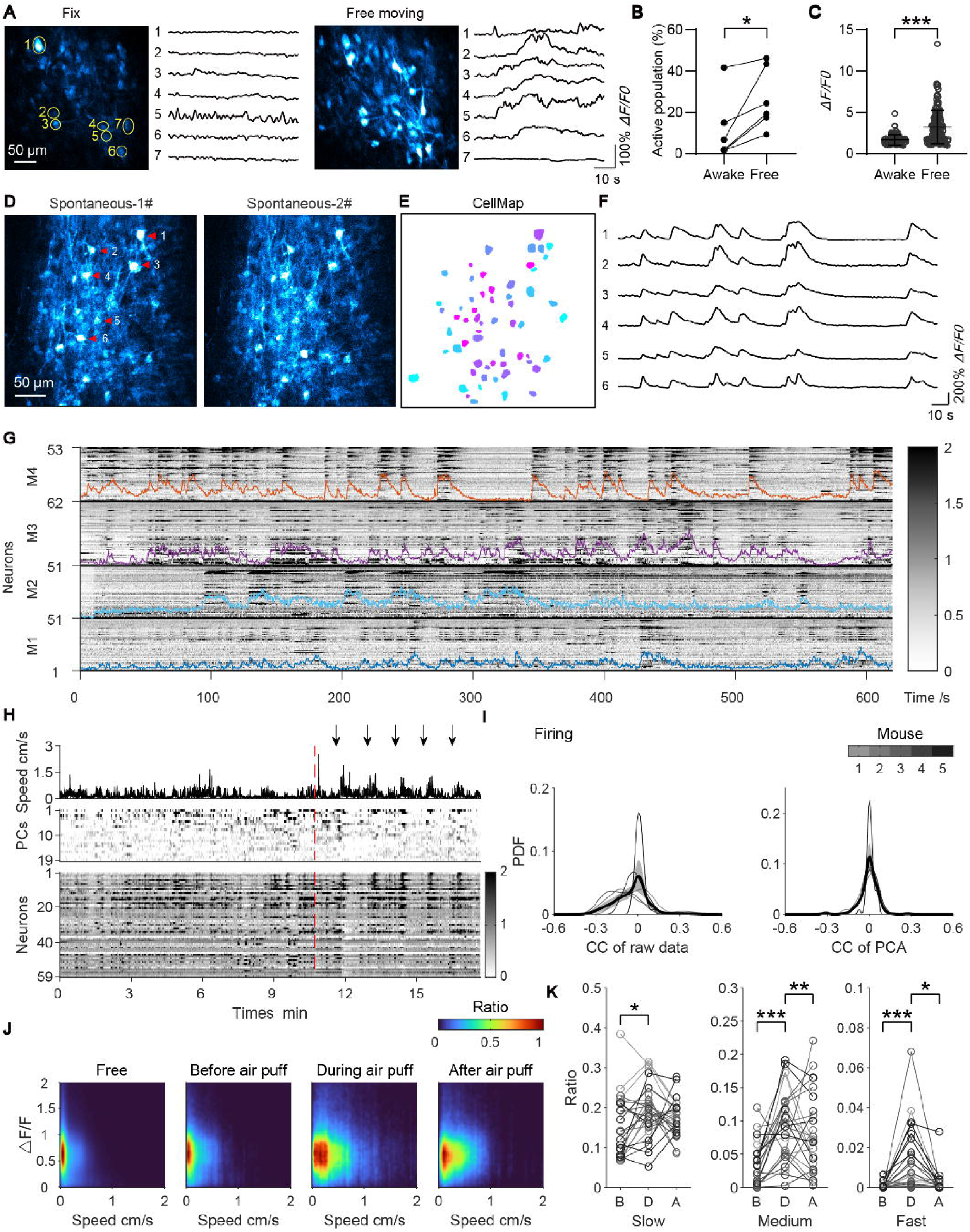
In vivo two-photon calcium imaging in the spinal cord. **A)** Representative two-photon in vivo image of the spinal cord in an awake fixed or free-moving state. Right: yellow circles represent representative identifiable neurons. Left: Ca^2+^ traces the same neurons over a 60 s period. **B)** The proportion of spontaneously active neurons in awake and free-moving states (N=6 mice per group; \*\**P*<0.05). **C)** The average maximum response amplitude of individual neurons, (Awake, N=51 neurons; free-moving, N=97 neurons; N=6 mice per group; \*\*\**P*<0.001). **D)** Representative two-photon image showing spontaneous activity of neurons in the spinal dorsal horn in a free-moving state. **E)** ROI contour map of an identified cell bodies. The purple to blue color gradient indicates the average brightness intensity of cell bodies from strong to weak. **F)** Ca^2+^ traces of neurons indicated in D. **G)** Heat maps showing the cellular activity of 4 representative mice (M1-M4) when they moved freely in a circular open field for 10 min. The left ordinate shows the number of cells extracted from each mouse, and the horizontal coordinate shows the recording time. In the spinal cord, there is a high synchronization of spontaneous neuronal activity. Curves of different colors represent the number of neuronal activity simultaneously. **H)** Speed (top), neural activity principal component (middle), raw neural activity data (bottom) from one mouse. Arrows indicate the onset time of the air puff (which lasted 5–10 s). PCA showed that 19 dimensions could explained 90% variance. With the red line as the boundary, the left side indicates the time when mice were moving freely without stimulation. Data showed that both the speed of the mice and the neural activity increased after the air puff (N=1 mice). **I)** Probability distribution of correlation coefficients between raw neural activity data and speed (left), and correlation coefficients between principal components of neural activity and speed (right). CC, Correlation Coefficient. The correlation coefficient indicated that both the original data and PCA data are independent of speed. (N=5 mice per group; each curve represents data from an individual animal) **J)** The heatmap of neural activities correlated with speed. The ratio at which a particular intensity of neural activity is generated at a particular speed. These particular intensities of neural activity were generated by specific behaviors (Free, before wind, during wind, after wind, respectively, data reflects 10 s of each). Note that the ratio increased during and after air puff stimuli. **K)** Neural activity rates at different speed levels (Slow: 0–0.25 cm/s, Medium: 0.25–1 cm/s, Fast: 1–2 cm/s). Note that the ratio is significantly higher during air puff stimuli (N=5 mice; each curve represents data from individual trials, 5 trials in 1 mouse; \**P* < 0.05; \*\**P* < 0.01; \*\*\**P* < 0.001). ROI, region of interest; PCA, principal component analysis.

### Calcium imaging in the spinal cord of free-moving mice with wind, pinch, or ice stimuli

To further probe dorsal horn neuron activity in awake mice, we measured noxious or non-noxious stimulus-evoked sensory neuron responses. Here, we used in vivo calcium imaging to investigate the activity of dorsal horn neurons in response to cutaneous stimuli induced by airpuff (wind), pinch, and 15 s or 90 s of ice stimulation. In this experiment, the same group of neurons was imaged before, during, and after each cutaneous stimulus. All stimulus modalities were applied successively in the same experiment, allowing us to assess the level of polymodality in this population of cells. We observed that cutaneous stimuli elicited robust Ca^2+^ transients in free-moving mice **(Fig. 3 A–B)**. We present the maximum response amplitudes of approximately 248 neurons responding to the wind, pinch, and ice stimuli, and sorted them based on 15 s of ice stimulation. Compared to 15 s ice stimuli, pinch or air-puff elicited significantly different activity in the same set of neurons, in contrast to 90 s ice stimuli, where neuronal activity was similar **(Fig. 3 C–D)**. In addition, pinch or air-puff did not evoke changes in the maximum response in maximum response amplitudes compared with ice **(Fig. 3 E)**. After cluster analysis, we found that the pinch-induced neuronal response was markedly different from that of other stimuli. In addition, the clustering results for ice-induced stimuli showed that neuronal responses were similar between different exposure times. The Venn diagram shows that 40% of cells responded to all three stimuli, whereas 16%, 10%, and 11% of cells responded to ice and wind, wind and pain, and pain and ice, respectively. The proportions of neurons, which responded to pain, ice, or wind alone were 8.6%, 8.1%, and 6.1%, respectively **(Fig. 3 F–G)**.

**Figure 3.**
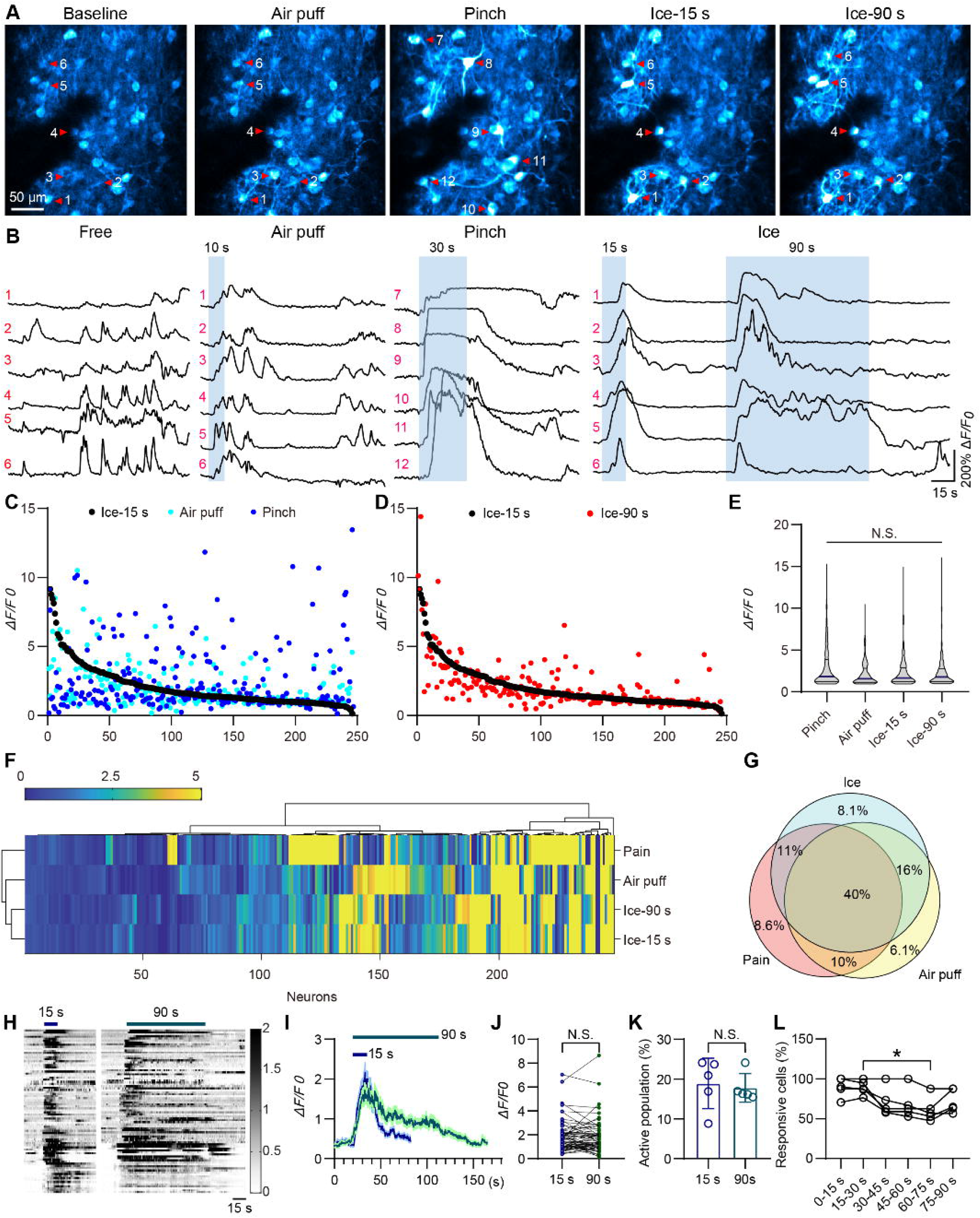
Sensory information encoding by dorsal horn neurons. **A)** Representative two-photon images of dorsal horn neurons expressing GCaMP6s in mice presented with various external stimuli (Air puff, Pinch, and Ice). Scale bar, 50 μm. **B)** Ca^2+^ traces of neurons indicated in A). **C–D)** A rank-ordered plot of maximum response amplitudes of all 248 neurons. Black, Ice-15 s; cyan, Air puff; blue, Pinch; red, Ice–90 s. **E)** *ΔF*/*F0* of all dorsal horn neurons following Pinch (N=192 neurons), Air puff (N=194 neurons), Ice–15 s (N=207 neurons), and Ice–90 s (N=191 neurons). Gray envelopes indicate the median (N=5 mice per group; *N.S. P* > 0.05). **F)** Cluster diagram of neuronal activity under different external stimuli (Air puff, Pinch, Ice). **G)** Venn plot shows the number of activated neurons in each group. **H)** Heat maps showing the activity of all 80 neurons that were activated after Ice stimuli. Each row in the heat map represents the response from the same neuron across the whole stimulus duration. Scale bar, 15 s. **I)** Population average *ΔF*/*F0* of all activated neurons following Ice–15 s or Ice–90 s stimuli. Blue and green lines indicate the mean value, and light green and light blue envelopes indicate the SEM. **J)** *ΔF*/*F0* of all sensory neurons following Ice–15 s, Ice–90 s (N=5 mice per group; *N.S. P* > 0.05). **K)** The percentage of neurons activated by different ice stimuli (N=5 mice per group; *N.S. P* > 0.05). **L)** The proportion of neurons activated in different time periods during Ice–90 s stimulation. Each curve represents one mouse. Note that during the late stage of ice stimulation, the proportion of neurons activated decreased significantly (N=5 mice per group; *N.S. P* > 0.05). *ΔF*/*F0*, maximum response amplitude; SEM, standard error of the mean.

In general, neurons which respond to cold stimuli respond robustly but rapidly adapt whilst under general anesthetic ^5^. The mechanisms underpinning this adaption occurring during free behavior remain unknown. We therefore investigated neuronal activation in response to ice stimuli of different exposure lengths in free-behaving mice. In response to short-term (15 s) and long-term (90 s) exposure to ice stimuli, we observed robust and reliable calcium transients in subsets of spinal dorsal horn neurons **(Fig. 3 A; Supplementary fig. 2)**. The heatmap shows calcium transients in neurons responding to cold throughout the whole stimulation period (**Fig. 3 H**), and the variability in responses further illustrates that neurons, which responded to cold stimuli responded and adapted quickly. These data suggest that the natural response to cold stimuli occurs in unique ensembles of spinal dorsal horn neurons. In addition, ice-evoked calcium transients reliably occurred at the onset of ice stimulation, and their duration was closely correlated with stimulus duration. A long ice stimulus further increased calcium response time **(Supplementary movie 2)** but did not evoke changes in the maximum response amplitudes or the proportion of the cell population activated (active population: 18.9±2.6% for 15 s stimulation versus 17.8±1.5% for 90 s stimulation, data shown as mean ± SEM) **(Fig. 3 I–K)**. These data suggest that cooling responses are mostly determined by the minimum temperature of the external stimulus. Despite responses to long-term ice stimulation gradually decreasing during the 90 s test, their adaptation to a sustained low temperature meant that the maximum response magnitude was determined by the absolute temperature, as the amplitudes decrease in response to a stable baseline temperature within just over a minute **(Fig. 3 L)**. These data demonstrate both sensitivity and adaptation of spinal neurons. Repeated administration of these temperature ramps (‘simple cooling ramps’) resulted in a small but non-significant decrease in the percentage of responsive cells **(Supplementary fig. 2 A–B)**. We recorded the stimulation temperature during 10 cooling trials, and the results of in vivo imaging showed that a similar group of neurons was activated in response to ice stimulation across all 10 trials in one mouse **(Supplementary fig. 2 C–D)**. On average, the maximum response amplitudes of cooling-evoked responses were similar and reliable, but varied within 7.03% of the mean across 10 trials **(Supplementary fig. 2 C–F)**. The sensitivity of responses was reliable, and the response periods stable in multiple mice **(Supplementary fig. 2 G)**.

### Long-term imaging in the spinal cord of free-moving mice

To quantify the clarity and stability of long-term two-photon imaging, we tracked the same cohort of axons of the spinal cord in Thy1-GFP mice over 2 months. Representative two-photon images of the spinal neuron axons were recorded at 1, 7, 14, 21, and 60 days, despite repeated placement and removal of the headpiece **(Fig. 4 A, Supplementary movie 3)**. We calculated the maximum cumulative projection of image displacements, which were generally around 10 μm in the mediolateral direction **(Fig. 4 B)**, and <25 μm in the rostrocaudal direction depending on the day **(Fig. 4 C)**. Interestingly, the image displacements at 60 days were significantly smaller in the mediolateral direction, which may indicate improved stability **(Fig. 4 B)**. To assess the potential impact of vertebral window implantation on the animals’ motility over an extended period, we assessed the cumulative time of moving distance and the average speed during a 5-min openfield test **(Fig. 4 D)**. We found that neither total distance nor average speed changed dramatically over 21 days, but both were significantly increased after 60 days **(Fig. 4 F–G)**. These results demonstrate the robustness of the system and its utility for repeated long-term in vivo imaging.

**Figure 4.**
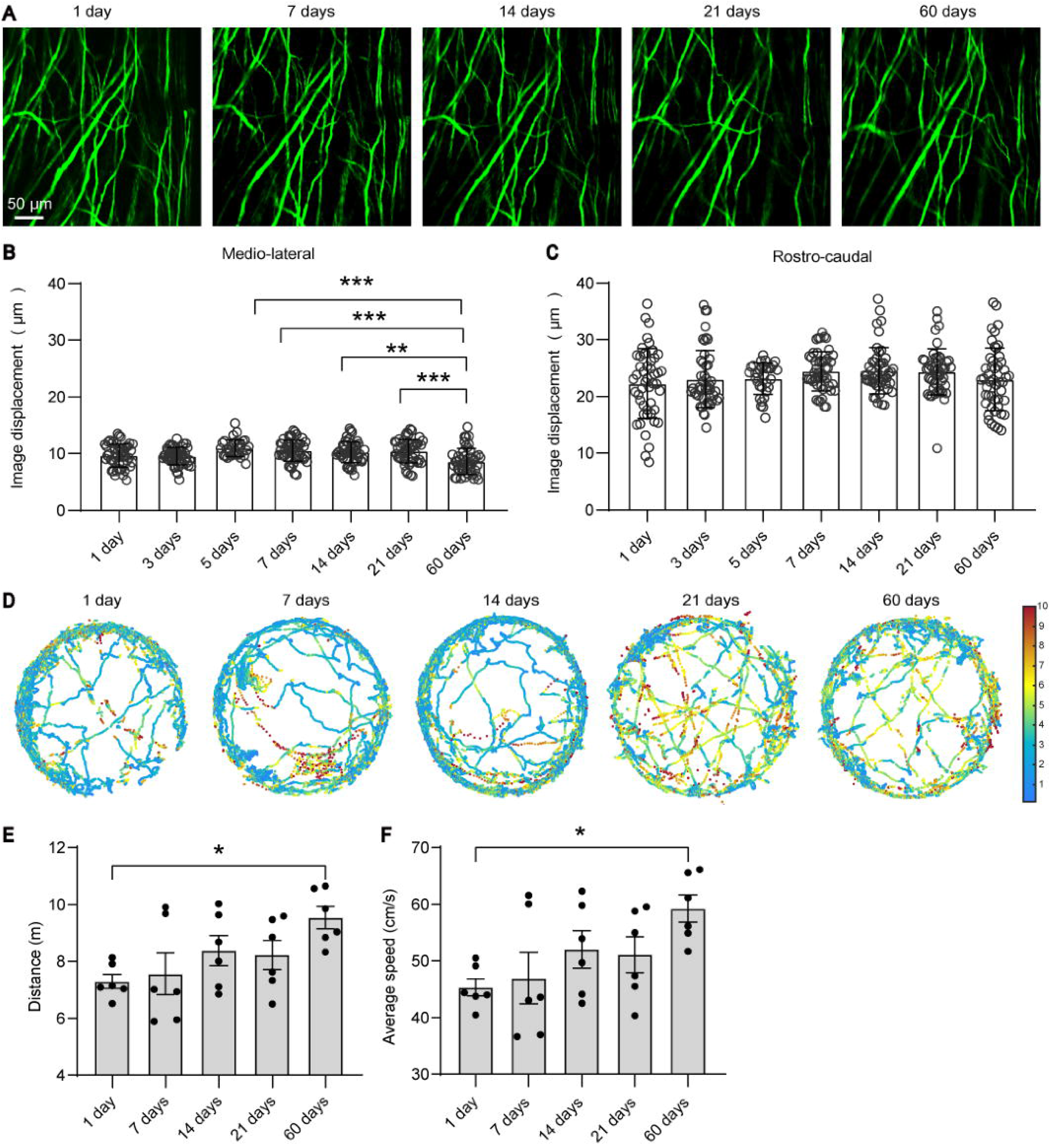
Long-term two-photon imaging of axons. **A)** Representative two-photon images of axons of spinal cord neurons expressing GFP. Scale bar, 50 μm. **B–C)** Population data showing image displacement in mediolateral and rostrocaudal directions occurring over 5 min. (N=6 mice per group; *N.S. P* > 0.05, \**P*<0.05, \*\**P*<0.01, \*\*\**P*<0.001). **D)** Heat map showing the motion track of mice in the circular open field over 60 days. The red to blue color gradient indicates the speed from strong to weak. **E)** Population data showing total distance and average speed in 5 min OFT for mice over 60 days (N=6 mice per group; \**P*< 0.05). GFP, green fluorescent protein; OFT, open field test.

Next, we performed in vivo Ca^2+^ imaging to assess the quality of the somatic activity of dorsal horn neurons. We injected AAV9-CaMKII-GCaMP6s at the lumbar level on the day of spinal cord window implantation and waited for 30 days for mouse recovery and viral expression. The images were recorded weekly for the next 21 days. We showed marked spontaneous Ca^2+^ transients of five individual neurons in one mouse **(Fig. 5 A)** and of the whole responsive populations in two mice **(Fig. 5 B)** during the 2 min open-field test. Next, to further investigate whether the functional plasticity of dorsal horn neurons changes in response to cutaneous stimuli due to the potential impact of vertebral window implantation on the animals’ back over an extended time period, we performed a 15 s ice stimulation trial and traced both individual neurons **(Fig. 5 C)** and associated subsets **(Fig. 5 D)** weekly. Three weeks of repeated administration of ice stimuli resulted in similar patterns of neuronal activity, which had a duration that was closely correlated with stimulus duration, and had no effects on the maximum response amplitude or total integrated calcium activity **(Fig. 5 E–F)**. These demonstrate stereotyped responses of dorsal horn neurons to cutaneous stimuli in healthy animals, which may contribute to the studies of the functions of the spinal cord in sensory perception.

**Figure 5.**
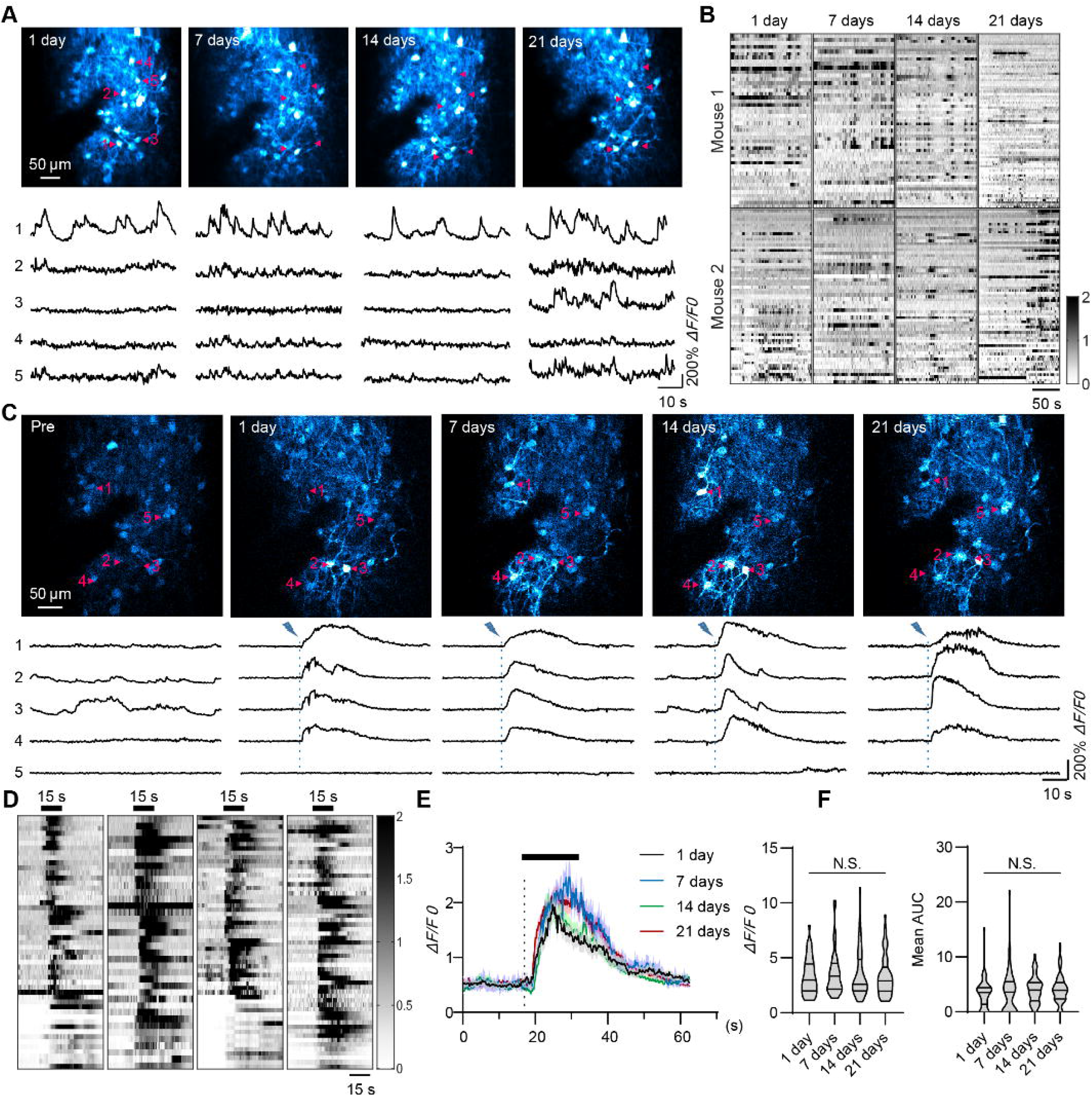
Long-term two-photon imaging of the activity of sensory neurons. **A)** Representative two-photon images and fluorescence traces of sensory neurons expressing GCaMP6s over 21 days. Scale bar, 50 μm. **B)** Representative heatmap showing the spontaneous activity of spinal dorsal horn neurons of 2 mice. Scale bar, 50 s. **C)** Representative two-photon images and fluorescence traces of sensory neurons expressing GCaMP6s, before and after ice stimulation. Scale bar, 50 μm. **D)** Heat maps showing the activity of all neurons that were activated after ice stimuli over 21 days (N=5 mice per group). Each row in the heat map represents the response from the same neuron throughout the whole stimulation duration. Scale bar, 15 s. **E)** Population average *ΔF*/*F0* of all activated neurons following ice stimulation over 21 days (N=5 mice per group). Black, 1 day; cyan, 7 days; green, 14 days; dark red, 21 days; light color envelope indicates SEM. Black line denotes the position of ice stimulating process. **F)** Maximum response amplitudes (left) and Integrated Ca^2+^ activity (right) of active neurons over time (N=5 mice per group; *N.S. P* > 0.05). *ΔF*/*F0*, maximum response amplitude; SEM, standard error of the mean.

### Calcium imaging following sciatic nerve injury

Neuropathic pain represents a common clinical disease ^21–23^. To further develop the imaging system and understand the response profiles of target spinal neurons in neuropathic pain, we applied the sciatic nerve injury (SNI) model and different stimuli to observe specific characteristics of neuronal sensitization in the dorsal horn such as hyperalgesia. Fourteen days after SNI surgery, we performed von Frey tests and found that the withdrawal threshold declined from 0.2 to 0.03 g **(Fig. 6 A)**. We also delivered 30 s pinch stimuli to freely moving mice and monitored the associated calcium transients in spinal neurons before and after SNI surgery. We observed sharp and instantaneous brightness changes at the moment of pinching **(Fig. 6 B)**. We further traced and analyzed the positive responsive neuron calcium release modes, and the entire response period lasted 150 s. We found that the maximum magnitude and active population of dorsal horn neurons were similar before and after SNI surgery **(Fig. 6 C;** *ΔF/F0*: 2.1±0.1 Pre versus 2.2±0.1 at 14 days after SNI surgery, data shown as mean ± SEM). The variation in responses to the pinch further illustrated quick neuronal responses and adaptation after SNI **(Fig. 6 D)**. However, neurons took longer to return to baseline after SNI compared to before **(Fig. 6 D)**, and the response time to pain increased significantly. Further analysis of the distribution showed that the difference mainly occurred 65–165 s after pinch stimuli (**Fig. 6 E;** where 0–15 s is pre-stimuli, 15–45 s is during pinch stimuli, and 45–165 s is during recovery). These data indicate that hyperpathia occurs in a chronic pain model of free-moving mice. Taken together, our results suggest that patterns of plasticity in spinal cord neurons changes in chronic disease states to adapt to the natural environment. In addition, spontaneous pain and allodynia are other characteristics of neuropathic pain, which are the most clinically problematic forms of pain ^23–25^. Although these pains are referred to the periphery, they arise within the CNS. Long-term sensitization of neurons suggests that increased calcium concentration in the dorsal horn is closely related to the central sensitization to neuropathic pain, and is thought to be part of many chronic pain sensitization processes ^2^. To study neuronal activity after SNI, we imaged the same population of sensory neurons in free-moving mice. We demonstrated that abnormal Ca^2+^ activity in the somas of sensory neurons continually appeared 14 days after SNI. In a 10-min open field test, transient spontaneous Ca^2+^ transients were still observed. However, a novel, usually pain-sensitive population of neurons became hyper-activated at 14 day after SNI. We named them ‘pain-sensing neurons’. Because pain-sensing neurons also undergo plastic changes in SNI model. When we referred the pain-sensing neurons appeared before SNI (Pre-pain), the percentage of neurons classified as pain-sensing neurons markedly increased from 28.21% in pre-to 60.96% in 14 days-SNI. When we referred the pain-sensing neurons appeared after SNI (SNI-pain), the percentage of neurons classified as pain-sensing neurons is also significantly higher than that in pre-SNI (61.96%). Meanwhile, we found that neurons undergo plastic changes in response to non-traumatic external stimuli in this model. Although the response distribution pattern and magnitude of individual cells remained unchanged throughout the stimulation period, the number of ice-activated cells but not air puff-activated cell increased significantly at 14 day after SNI**. (Supplementary fig. 6)**. However, non-noxious stimuli also induced hyper-activated neurons which usually is pain-sensitive after SNI. When we referred the pain-sensing neurons appeared before SNI, the percentage of neurons classified as pain-sensing neurons rose from 26.42% in pre-to 57.62% in 14 days-SNI after air puff stimuli, and rose from 35.78% in pre-to 78.40% in 14 days-SNI after ice stimuli. When we referred the pain-sensing neurons appeared after SNI, the percentage of neurons classified as pain-sensing neurons is also significantly higher than that in pre-SNI (air puff stimuli: 63.05%; ice stimuli: 71.04%).

**Figure 6.**
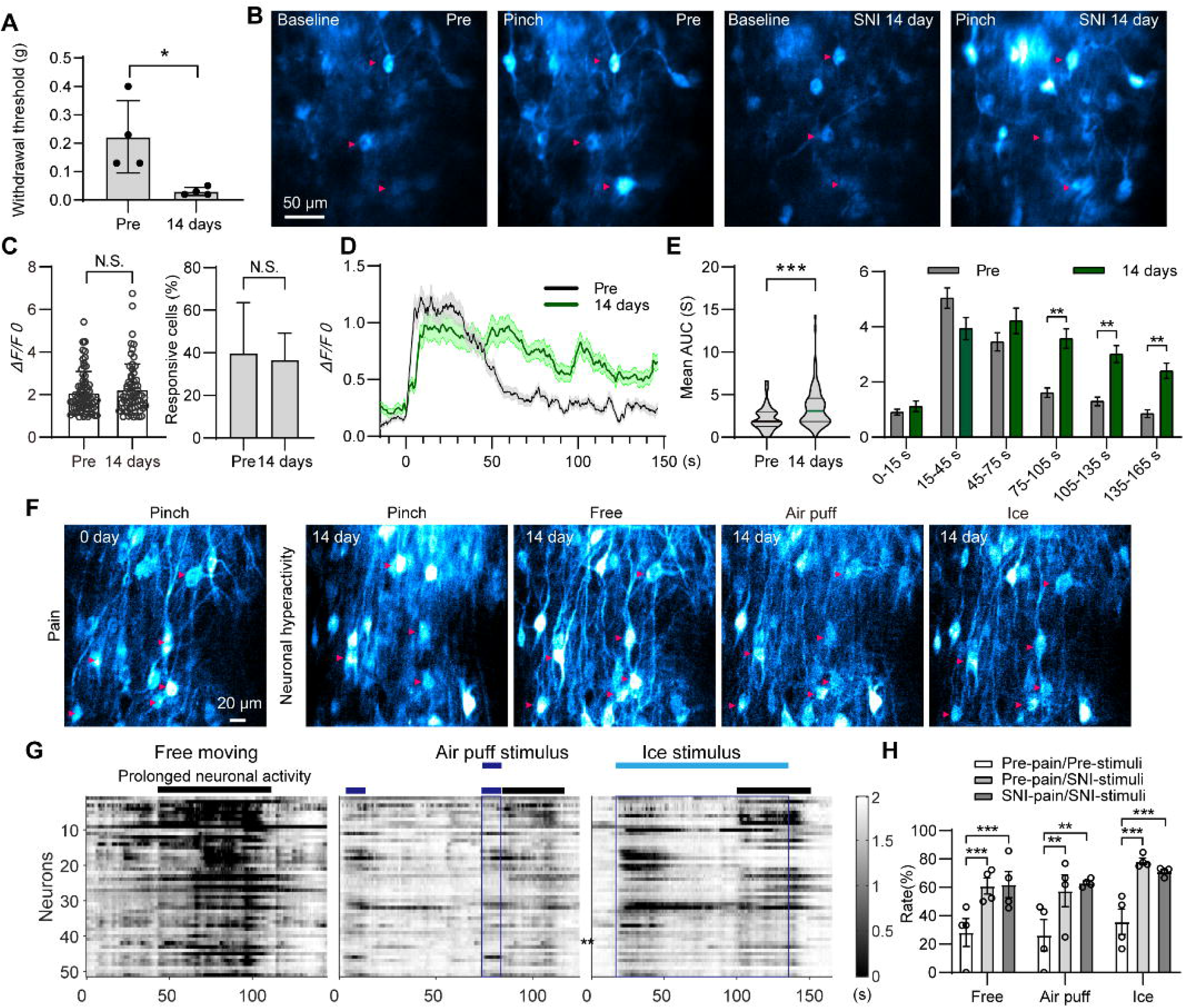
Sciatic nerve injury induces long-lasting increases in neuronal Ca^2+^ activity in neuropathic pain. **A)** Withdrawal threshold test (N=4 mice per group; \**P*<0.05). **B)** Representative two-photon images of the response of dorsal horn neurons to pinch stimulation before and after SNI. Scale bar, 50 μm. **C)** Left: *ΔF*/*F0* (maximum response amplitudes) of all sensory neurons during pinch stimuli. Right: the percentage of neurons activated by pinch stimuli before and after SNI (N=4 mice per group; *N.S. P* > 0.05). **D)** Population average *ΔF*/*F0* of all activated neurons following pinch stimuli before and after SNI. Black, Pre; green, 14 days after SNI. The light color envelope indicates SEM. **E)** Total integrated Ca^2+^ activity of active neurons before and 14 days after SNI (left). The integrated Ca^2+^ activity in different time periods during pinch stimulation (right). Note that during the late stage of pinch stimuli, the integrated Ca^2+^ activity activated is significantly prolonged 14 days after SNI (N=4 mice per group; \**P*<0.05, \*\**P*<0.01, \*\*\**P*<0.001). **F)** Representative two-photon images of the responses of sensory neurons to pinch stimuli before SNI and images of abnormal neuronal responses to various external stimuli after SNI. Scale bar, 50 μm. **G)** Heat maps reflecting the activities of the abnormal neuronal responses to different external stimuli in the same FOV (an example field of view) after SNI (N=1 mice). **H)** Percentages of abnormal neuronal response exhibiting Ca^2+^ transients during different external stimuli compared with pain before and after SNI (N=4 mice per group; \*\**P*<0.01, \*\*\**P*<0.001). SNI, sciatic nerve injury; *ΔF*/*F0*, maximum response amplitude; SEM, standard error of the mean; FOV, field of view.

## Discussion

As a bridge connecting the brain and the peripheral nervous system, the spinal cord transmits neural information to coordinate normal activities and plays an important role in somatosensory and motor processes ^1,26–28^. Recently, with the combination of miniaturized microscopy and calcium imaging technology, it can acquire images in freely moving animals ^17,18^. Application of this technology has spurred on a certain degree of creativity and innovation in the field of brain imaging, but remains challenging for spinal cord imaging. The complex structure, vigorous and irregular tissue movement, long-term imaging instability, and the difficulty of stable implantation of a miniaturized microscope have led to a series of challenges in spinal cord intravital imaging, such as focal plane drift and image distortion or loss ^13,14,29^. Although the pioneering of miniaturized one-photon microscopy has allowed spinal imaging in freely behaving mice, the restrictive calcium imaging period has excluded the possibility for long-term investigation, and results may have been affected by severe inflammation ^19,30,31^. Moreover, the principles of miniaturized one-photon microscopy imaging limit the achievable contrast and are restricted by differences in image contrast between the white and gray matter of the spinal cord ^13^. We developed a new intervertebral fusion method by clamping adjacent vertebral bodies with two custom-designed stainless steel grooves to minimize spinal movement during imaging. The method developed in our study allows the monitoring of responses to cutaneous stimuli in the dorsal horn neurons of the spinal cord of freely behaving mice over minutes to weeks. Our most prominent improvement lies in spinal fixation. Image artifacts caused by motion can be further reduced by gently pressing the coverslip during image collection. Using this method, in vivo imaging of the spinal cord of freely behaving mice can be readily achieved without significant effects on the animals’ motor or sensory functions. Following window implantation, the mouse motor ability is inevitably reduced. However, it does not affect the basic motor function and sensation in mice, which is consistent with previous studies ^15,19,20^. Our method of spinal fixation dramatically decreases the loss of imaging artifacts by reducing irregular movement of the spinal cord, and it can obtain images of stable cellular activity from the spinal cord of freely behaving mice using miniaturized two-photon microscopy.

A core challenge of neurobiology remains to understand the connection between sensory transduction in the periphery and associated behavior ^32–34^. Several studies have shown that neurons in lamina I are mainly sensory neurons ^9,12,35^, but whether sensation is affected by locomotion or is responsive to stimulation remains unclear. Our results show that differences in perception in response to external environmental stimulation are not due to mouse speed. This may indicate that responses of layer I neurons which are part of the ascending pathway mainly contribute to perception rather than locomotion. Neurons in lamina I are mainly classified into nociceptive-specific cells, multi-type nociceptive cells, and temperature-specific cells ^6,36,37^. Our results also show that there are both separate and overlapping subpopulations of neurons that respond to pinch, air puff, or cold stimuli. Electrophysiological and one-photon microscopy imaging studies have shown that different sub-modal skin perceptions can induce overlapping neuronal responses during anesthesia and free movement ^4,19^. Nevertheless, we found that subsets of responsive neurons in lamina I overlapped, and that firing rates were higher than previous results obtained using one-photon imaging ^19^. There are two potential reasons, which may account for this phenomenon. The first is that interference of background fluorescence during one-photon imaging may lead to difficulties in signal identification. The second is that recording usually took place 30 days after device implantation, which reduced the effects of background noise and inflammation ^30^. The spinal dorsal horn neurons display an inherent stereotyped firing pattern in response to external stimuli ^4,8^. Our results show that the intensity of neuronal activity is due to stimulus duration as well as stimulus type. A cluster analysis demonstrated that the distribution of neural calcium responses induced by noxious and non-noxious stimuli was different and encoded at the single-neuron level. These results indicate that spinal dorsal horn neurons have an innate coding mode for external nociceptive and non-noxious stimuli ^6^. In addition, after prolonging the duration of ice stimulation, we found that the distribution of neural calcium responses induced by ice stimulation was similar. These results indicate that spinal dorsal horn neurons can quickly adjust to external environmental stimuli at the single-cell level by encoding specific cell groups.

Neuropathic pain represents a common clinical disease ^21–23^. Peripheral nerve disorders can cause chronic pain, such as diabetes, uremia, neuralgia after herpes zoster, and ischemic neuropathy ^38–41^. However, the pathogenesis, disease progression, and the effectiveness and appropriate duration of treatments remain unclear and difficult to assess ^2,12,24,26,38^. By imaging spinal cord axons over 60 days, we have shown the strong stability of our system and demonstrated minimal deformation and displacement with our custom-designed components on imaging clarity and mouse behavior. With this system, we are able to follow the activity of spinal cord sensory neurons during disease progression. In the case of neuropathology, the central mechanism mainly includes the sensitization of dorsal horn neurons, especially enhanced response to noxious stimuli (hyperalgesia), abnormal response to non-noxious stimuli (allodynia), and spontaneous pain caused by increases in spontaneous impulses ^2,3,42–44^. A previous study showed that the sensitization of dorsal horn neurons is closely related to excitatory amino acids, NMDA and AMPA receptors, substance P, intracellular calcium ions, and protein kinases, though we still lack data regarding mechanisms at the cellular level ^2,45,46^.

We induced neuropathic pain using a SNI model and monitored both the preoperative and postoperative Ca^2+^ activity of sensory neurons. By monitoring the activity of individual neurons in the spinal cord of free-moving mice over weeks, we demonstrate that responses of dorsal horn neurons are stereotyped in healthy mice, but that hyperactivity and functional plasticity occur in mice with neuropathic pain in free-behaving states. Our results showed that hyperalgesia is mainly caused by abnormally enhanced firing of nociceptive neurons. More importantly, abnormal prolonged Ca^2+^ activity of nociceptive neurons induced by non-noxious stimuli appeared after SNI but rarely appeared in healthy mice, which provide some insight for mechanism for allodynia and spontaneous pain. These results are consistent with previous findings from electrophysiology research ^24^. In addition, we found that in the neuropathological model, the response of dorsal horn neurons to certain non-noxious stimuli undergoes changes in plasticity, which mainly manifests as changes in the active neuronal subsets, while inherent patterns of single cell firing remained the same.

In summary, we have innovated the application of miniaturized two-photon in vivo imaging to the spinal cord of mice, which can help us study the encoding of sensory inputs and motion in freely moving situations. The mechanical design and surgical preparations facilitate optical access to the mouse spinal cord. This means it is possible to extend the application of miniaturized two-photon microscopy to the study of the neuronal dynamics of cutaneous stimuli such as the coding of cutaneous temperature, air puff, and pinch in the spinal cord, and particularly in chronic processes such as neuropathic pain. Combining in vivo imaging with animal behavior will help unravel maladaptive changes in the spinal circuitry and provide therapeutic insights into these devastating disorders.

## Methods

### Experimental animals

C57BL/6 wild-type mice (8–10 postnatal weeks) and transgenic mice expressing green fluorescent protein in layer 5 pyramidal neurons (Thy1-GFP line M) were purchased from the Jackson Laboratory. All animals were housed under a 12/12-h light/dark cycle at 22°C and had free access to clean food and water. Mice were housed individually to minimize the risk of injury after surgery. All procedures were approved by the Animal Care and Use Committee of the Shenzhen Institute of Advanced Technology (SIAT), Chinese Academy of Sciences (CAS).

### Surgical procedures for spinal window implantation

Each mouse was anesthetized by inhalation of 1.5–3% isoflurane (RWD Life Science, Shenzhen, China) in 100% O2 and placed on a stereotactic stage (RWD Life Science, Shenzhen, China). We performed a dorsal longitudinal incision and bluntly dissected connective tissue and muscle covering the vertebrae from L3 to L6 **(Fig 1 B; Supplement fig 1 B)**. After carefully cleaning the muscle, blood, and membrane tissue attached to the spinous processes and both transverse processes of the spine and surrounding bone surfaces (L3 to L6 vertebrae), we inserted a stainless steel groove into the lower part of the vertebral transverse process in a stable position. To minimize motion artifacts, the entire transverse process was placed over the stainless steel groove on both sides, and both the stainless steel grooves were adjusted using stereotaxic apparatus until the wobble of the vertebrae had been suitably reduced **(Supplement fig 1 A–B)**. We then cleaned the field with H_2_O_2_ for hemostasis and debridement and dropped 3M Vetbond Tissue Adhesive (Amazon) to prevent tissue effusion. We gently removed the raised part of the vertebral transverse process around L4 or L5 with a high-speed micro-drill (tip diameter 0.5 mm), leaving the surface of the trimmed bone and spinal cord in the same optical plane. After gentle washing Physiological saline, and when the bleeding had stopped, we had access to a clean window with an intact dura mater. After the surface of the spinal cord was cleaned with sterile saline, we dropped a thin layer of Kwik-Sil^®^ silicone elastomer to the surface and covered the window with a 4–5 mm coverslip with minimal pressure until the diameter of the posterior artery was reduced by approximately 50%. We gently sucked out the effusion and added 3M glue while maintaining the pressure. Two minutes later, we sealed the edges and glued the skin to the chamber with cyanoacrylate adhesive (fast curing super glue 406) and then dental cement. We continuously delivered intraperitoneal injections of penicillin (50000 units/kg) mixed with dexamethasone (0.2 mg/kg) for 7 days and tramadol (0.3 ml/kg) for 7 days postoperatively.

GCaMP6s was expressed with a recombinant adeno-associated virus (AAV) under the calmodulin-dependent kinase II promoter (2.26×10^12^ GC per ml). After exposure of the L3 or L4 spinal cord segment, we placed a piece of hemostatic sponge to the surface to keep it moist and clean. Then, 0.3 μl of AAV9-CaMKII-GCaMP6s was injected manually within 10 min 300–400 μm left of the center of the posterior artery (at a depth of 150–200 mm below the dura) with an injector (Thinker Tech) and glass microelectrodes. Next, we added a drop of Kwik-Sil^®^ silicone elastomer to the target region. Ten minutes later, we carefully lifted the microelectrodes and moved the silicone elastomer away. We then finished the process described above and kept the mouse warm until recovery.

### SNI model

SNI has been used to induce neuropathic pain as described in previous studies ^47–49^. Briefly, each mouse was deeply anesthetized by inhalation of 1.5–3% isoflurane in 100% O_2_ and placed on a stereotactic stage, and a longitudinal skin incision was carefully made after sterilization with 70% ethanol. Using forceps, the biceps femoris muscle fibers were separated to allow visualization of the sciatic nerve and its three terminal branches: the sural, common peroneal, and tibial nerves. SNI was induced by tying a tight ligature made from 5.0 silk sutures (RockTechtl) around the common peroneal and tibial nerves, and then transected distal to the ligation, removing approximately 4 mm of the distal nerve stump, whilst leaving the tibial nerve intact. The skin was then closed and stitched with 5.0 silk sutures (RockTechtl).

### Von-Frey test

Mechanical allodynia was measured using the top-down method ^50^ using von Frey filaments (Stoelting, Wood Dale, IL, USA), with ascending forces expressed in grams (0.02 to 2.0 g). Mice were habituated for 20–30 min in silent enclosures with a wire mesh floor. Each filament was applied five times in a row against the medial plantar surface of the hind paw. Prolonged hind paw withdrawal, rapid fanning, or licking induced by the filament was defined as a positive nociceptive response. If a positive response was observed, filament lower strength filament was applied, and if there was no response, a stronger filament was applied. The lowest gram-force eliciting positive responses in three of the five repeated stimuli was defined as the 50% withdrawal threshold. The pattern of responses was used to calculate the 50% threshold = (10[χ+κδ])/10,000), where χ is the log of the final von Frey filament used, κ is the tabular value for the pattern of responses, and δ is the mean difference between filaments used in log units. The log of the 50% threshold was used to calculate the summary and test statistics, in accordance with Weber’s law.

### Freely moving mouse two-photon imaging

We used a compatible scan lens to support the FOV for imaging. The excitation wavelength was determined to be 920 nm. Imaging FOV: 420×420 μm, XY pixels: 512×512, acquisition frame rate: 4.84 Hz. Thirty days postoperatively, the mice were assessed and used for imaging. With a clean window and intact motor ability, the mouse was restrained with a special holder and the 2.5×2.5 mm^2^ square window over the target region of dorsal horn was targeted with FHIRM-TPM. A well-matched holder was mounted in the chamber and reinforced with a cyanoacrylate adhesive. A detachable 2.8 g lens with fiber and cable was fixed to the assembly for imaging. Individual mice carrying the assembly with or without lenses were tested in the behavioral experiments.

### Behavioral tests

The mice were subjected to an open field test. Mice were acclimatized to the environment for 1–2 min before the test and placed in a circular open field (diameter 50 cm, height 50 cm). Each mouse was allowed to rest before trials, and all videos were simultaneously recorded with a camera from the top of the cage. Motion test: mice with FHIRM-TPM were placed in the testing field and moved freely for 10 min. Air puff: the mouse was placed in the field. Compressed air was used to stimulate the mouse with acute airflow, and the mouse stimulation was delivered continuously for 10 s. We recorded the free behavior and images of spinal cord neurons simultaneously. The stimulation was performed 5–6 times with an interval of approximately 90 s. Pinching: we used a custom-made clip (P=340 g) to clamp the whole hind paw or the medial plantar surface of the hind paw of the mouse for 30 s. Then, we removed the clip and allowed it to move freely. Ice treatment: a rat breeding box with ice was placed in the middle of the circular open field. The mouse was placed on ice for 15 or 90 s during recording. We repeated the 15 s ice trail 5–10 times between intervals of approximately 90 s.

### Data processing and analyses

Image data analysis: Based on synchronously recorded behavioral videos, we selected axon images in the target period and analyzed them with ImageJ. The specific steps were as follows: Use ‘Image’ > ‘Z-projector’ > ‘max intensity’ to obtain the maximum projection of the continuous image in the target time period. To compare the image displacement in different periods, we selected eight axon sites in the horizontal plane for displacement statistics. The average displacement value of the eight sites was taken as the index for motion artifacts.

Similarity assessment of spinal cord video frames: The MS-SSIM index was used to quantify the similarity between two adjacent frames of the spinal cord video. The ‘multissim’ function in MATLAB was used to calculate the MS-SSIM. The one-hundredth frame in each spinal cord video was selected as the reference image to calculate the MS-SSIM with each other frame in the same video. Then, the distributions of the MS-SSIM of each video were binned and plotted using 30 bars.

Calcium imaging data analysis: In this study, the somas of spinal cord neurons were labeled with GCaMP6s in the cytoplasm. Raw calcium imaging data were processed using customized software for non-rigid motion correction of calcium imaging data (LeapBrain RAYGEN HEALTH). Regions of interest (ROIs) corresponding to visually identifiable somas were selected for quantification of dorsal horn neurons. The fluorescence time course in each soma was measured using ImageJ by averaging all the pixels within the ROI covering the soma. *ΔF*/*F0* was calculated as *ΔF*/*F0* = (*F*–*F0*)/*F0*×100%, where F0 is the baseline fluorescence signal averaged over a 2–5 s (free-moving period corresponding to the lowest fluorescence signal over the recording period. Stimuli such as air puff, ice, and pinch were delivered, *F0* was the baseline fluorescence signal averaged over a 15–20 s period corresponding to the lowest fluorescence signal over the recording period. Active spinal cord neurons were defined based on the criteria that calcium transients were beyond the threshold of one times the standard deviation of the baseline. To compare the neuron release patterns under different stimuli and at different times, we counted the *ΔF*/*F0* during calcium release. To compare neuronal activity among different cells and at different time points, we performed an integrated measurement of a cell’s output activity over the total stimulus duration, termed total integrated calcium activity, and calculated the area under the curve (AUC). The total integrated calcium activity was the average of *ΔF*/*F0* over the total stimulus duration. The AUC is calculated as:

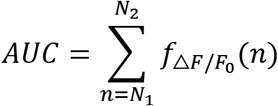

where *n* is the sample time point of calcium activity, *N*_1_ is the start time point, *N*_2_ is the end time point and *f*Δ*F*/*F*_0_(*n*) is the *ΔF*/*F0* of calcium activity.

PCA of neuron data: Using PCA, the best linear combinations of neural activities were extracted as principal components (PCs) to represent the main information. The ‘pca’ function in MATLAB was used with the dimension of neural activities considered as a variable. The first 19 PCs could explain more than 90% of the total variance, which were sorted by the eigenvalues and plotted.

Correlation analysis of speed and neural activities: The ‘jbtest’ function in MATLAB was used to test the normality of data before temporal correlation analysis. The ‘corr’ function in MATLAB was used to calculate the temporal correlations between speed and each neuronal activity or PC of mice. If the data followed a normal distribution, Pearson’s correlation was used. If the data were not normally distributed, Spearman’s correlation was used. The distribution of CCs was estimated using kernel density estimation. Because of the cluster gram of neurons with different stimulations, the peak values of each neuron with different stimulation periods were first arranged in a matrix. Second, hierarchical clustering was separately used for stimulation and neuron dimensions. Third, the standardized Euclidean distance was used to calculate the pairwise distance, and the inner squared distance was used for the clustering trees. Finally, according to clustering trees, neural activity data were sorted and plotted with dendrograms of stimulation modes and neuron dimensions.

Behavioral data analysis: Behavioral video clips of the mice were analyzed using the Anymaze animal behavioral analysis software and MATLAB. Through multi-view image tracking and track reconstruction, the movement distance and movement speed of the mouse during the analysis period were extracted as indices of the motion ability.

### Statistical analyses

Data were analyzed using GraphPad Prism 9.0 software. Before hypothesis testing, data were first tested for normality using the Shapiro-Wilk normality test and then for homoscedasticity using the F test. If the null hypothesis was that the data had a normal distribution and homogeneity of variances could not be rejected, parametric tests were used (for example, Student’s t-test for two groups, one-way ANOVA with Tukey’s multiple comparisons test for more than two groups). If the null hypothesis was that the data did not have a normal distribution and homogeneity of variances could be rejected, nonparametric tests were used (for example, Mann-Whitney test for two groups, one-way ANOVA with the Kruskal-Wallis test for more than two groups). NS, single, double, and triple asterisks indicate *N.S. P* > 0.05, * *P* < 0.05, ** *P* < 0.01, and *** *P* < 0.001, respectively.

## Acknowledgments

We would like to thank Peking University-Nanjing Joint Institute of Translational Medicine for supplying the LeapBrain to process the data. This work was supported in part by the National Key R&D Program of China (2018YFA0701403 to P.W.), Key Area R&D Program of Guangdong Province (2018B030338001 to P.W., 2018B030331001 to L.W.), the National Natural Science Foundation of China (NSFC 31500861 to P.W., NSFC 31630031 to L.W., NSFC 91732304 to L.W., NSFC 31930047 to L.W.), the Chang Jiang Scholars Program and the Ten Thousand Talent Program (to L.W.), the International Big Science Program Cultivating Project of CAS (172644KYS820170004 to L.W.), the Strategic Priority Research Program of Chinese Academy of Science (XDB32030100 to L.W.), the Youth Innovation Promotion Association of the Chinese Academy of Sciences (2017413 to P.W.), the Shenzhen Government Basic Research Grants (JCYJ20170411140807570 to P.W., JCYJ20170413164535041 to L.W.), the Science, Technology and Innovation Commission of Shenzhen Municipality (JCYJ20160429185235132 to K.H.), the Helmholtz-CAS joint research grant (GJHZ1508 to L.W.), the Guangdong Provincial Key Laboratory of Brain Connectome and Behavior (2017B030301017 to L.W.), the Guangdong Special Support Program (to L.W.), the Key Laboratory of CAS (2019DP173024 to L.W.), the National Natural Science Foundation of China (Grant No.81901984), the Shenzhen Key Science and Technology Infrastructure Planning Project (ZDKJ20190204002 to L.W.).

## Author contributions

F.J., T.-W.H, P.-F.W., and L.-P.W. designed the experiments. F.J. designed the vertebral mount, F. J. and K.J performed the experiments, Y.-N.H, W.-L.J, K.J, N.L and S.-Y.C analyzed the data. T.-W.H, Z.-H.L, P.-F.W., and L.-P.W. contributed to data interpretation. F.J., W.-L.J, Y.-N.H, T.-W.H, P.-F.W., and L.-P.W. wrote the manuscript.

## Competing interests

Correspondence to Pengfei Wei, Liping Wang.

## Ethics declarations

### Competing interests

The authors declare no competing interests.

## Notes

### Competing Interest Statement

The authors have declared no competing interest.

## References

1 Moehring, F., Halder, P., Seal, R. P. & Stucky, C. L. Uncovering the Cells and Circuits of Touch in Normal and Pathological Settings. Neuron 100, 349–360, doi:10.1016/j.neuron.2018.10.019 (2018).

2 Ji, R. R., Kohno, T., Moore, K. A. & Woolf, C. J. Central sensitization and LTP: do pain and memory share similar mechanisms? Trends Neurosci 26, 696–705, doi:10.1016/j.tins.2003.09.017 (2003).

3 Sandkuhler, J. Central sensitization versus synaptic long-term potentiation (LTP): a critical comment. J Pain 11, 798–800, doi:10.1016/j.jpain.2010.05.002 (2010).

4 Abraira, V. E. et al. The Cellular and Synaptic Architecture of the Mechanosensory Dorsal Horn. Cell 168, 295–310 e219, doi:10.1016/j.cell.2016.12.010 (2017).

5 Chisholm, K. I. et al. Encoding of cutaneous stimuli by lamina I projection neurons. Pain 162, 2405–2417, doi:10.1097/j.pain.0000000000002226 (2021).

6 Koch, S. C., Acton, D. & Goulding, M. Spinal Circuits for Touch, Pain, and Itch. Annu Rev Physiol 80, 189–217, doi:10.1146/annurev-physiol-022516-034303 (2018).

7 Cheng, L. et al. Identification of spinal circuits involved in touch-evoked dynamic mechanical pain. Nat Neurosci 20, 804–814, doi:10.1038/nn.4549 (2017).

8 Ran, C., Hoon, M. A. & Chen, X. The coding of cutaneous temperature in the spinal cord. Nat Neurosci 19, 1201–1209, doi:10.1038/nn.4350 (2016).

9 Haring, M. et al. Neuronal atlas of the dorsal horn defines its architecture and links sensory input to transcriptional cell types. Nat Neurosci 21, 869–880, doi:10.1038/s41593-018-0141-1 (2018).

10 Grudt, T. J. & Perl, E. R. Correlations between neuronal morphology and electrophysiological features in the rodent superficial dorsal horn. J Physiol-London 540, 189–207, doi:10.1113/jphysiol.2001.012890 (2002).

11 Peirs, C. et al. Mechanical Allodynia Circuitry in the Dorsal Horn Is Defined by the Nature of the Injury. Neuron 109, doi:10.1016/j.neuron.2020.10.027 (2021).

12 Todd, A. J. Neuronal circuitry for pain processing in the dorsal horn. Nat Rev Neurosci 11, 823–836, doi:10.1038/nrn2947 (2010).

13 Cheng, Y. T., Lett, K. M. & Schaffer, C. B. Surgical preparations, labeling strategies, and optical techniques for cell-resolved, in vivo imaging in the mouse spinal cord. Exp Neurol 318, 192–204, doi:10.1016/j.expneurol.2019.05.010 (2019).

14 Nelson, N. A., Wang, X., Cook, D., Carey, E. M. & Nimmerjahn, A. Imaging spinal cord activity in behaving animals. Exp Neurol 320, 112974, doi:10.1016/j.expneurol.2019.112974 (2019).

15 Chen, C. et al. Long-term imaging of dorsal root ganglia in awake behaving mice. Nat Commun 10, 3087, doi:10.1038/s41467-019-11158-0 (2019).

16 Huang, T. W. et al. Identifying the pathways required for coping behaviours associated with sustained pain. Nature 565, 86–+, doi:10.1038/s41586-018-0793-8 (2019).

17 Zong, W. J. et al. Miniature two-photon microscopy for enlarged field-of-view, multi-plane and long-term brain imaging. Nat Methods 18, 46–+, doi:10.1038/s41592-020-01024-z (2021).

18 Zong, W. J. et al. Fast high-resolution miniature two-photon microscopy for brain imaging in freely behaving mice (vol 14, pg 713, 2017). Nat Methods 14(2017).

19 Sekiguchi, K. J. et al. Imaging large-scale cellular activity in spinal cord of freely behaving mice. Nat Commun 7, 11450, doi:10.1038/ncomms11450 (2016).

20 Farrar, M. J. et al. Chronic in vivo imaging in the mouse spinal cord using an implanted chamber. Nat Methods 9, 297–U113, doi:10.1038/Nmeth.1856 (2012).

21 Aasvang, E. K., Brandsborg, B., Christensen, B., Jensen, T. S. & Kehlet, H. Neurophysiological characterization of postherniotomy pain. Pain 137, 173–181, doi:10.1016/j.pain.2007.09.026 (2008).

22 Rasmussen, P. V., Sindrup, S. H., Jensen, T. S. & Bach, F. W. Symptoms and signs in patients with suspected neuropathic pain. Pain 110, 461–469, doi:10.1016/j.pain.2004.04.034 (2004).

23 Sandkuhler, J. Models and Mechanisms of Hyperalgesia and Allodynia. Physiol Rev 89, 707–758, doi:10.1152/physrev.00025.2008 (2009).

24 Campbell, J. N. & Meyer, R. A. Mechanisms of neuropathic pain. Neuron 52, 77–92, doi:10.1016/j.neuron.2006.09.021 (2006).

25 Chao, D. et al. Dorsal root ganglion stimulation of injured sensory neurons in rats rapidly eliminates their spontaneous activity and relieves spontaneous pain. Pain, doi:10.1097/j.pain.0000000000002284 (2021).

26 Basbaum, A. I., Bautista, D. M., Scherrer, G. & Julius, D. Cellular and molecular mechanisms of pain. Cell 139, 267–284, doi:10.1016/j.cell.2009.09.028 (2009).

27 Bourane, S. et al. Gate control of mechanical itch by a subpopulation of spinal cord interneurons. Science 350, 550–554, doi:10.1126/science.aac8653 (2015).

28 Grillner, S. Biological pattern generation: The cellular and computational logic of networks in motion. Neuron 52, 751–766, doi:10.1016/j.neuron.2006.11.008 (2006).

29 Farrar, M. J., Rubin, J. D., Diago, D. M. & Schaffer, C. B. Characterization of blood flow in the mouse dorsal spinal venous system before and after dorsal spinal vein occlusion. J Cerebr Blood F Met 35, 667–675, doi:10.1038/jcbfm.2014.244 (2015).

30 Sommer, C., Leinders, M. & Uceyler, N. Inflammation in the pathophysiology of neuropathic pain. Pain 159, 595–602, doi:10.1097/j.pain.0000000000001122 (2018).

31 Inoue, K. & Tsuda, M. Microglia in neuropathic pain: cellular and molecular mechanisms and therapeutic potential. Nat Rev Neurosci 19, 138–152, doi:10.1038/nrn.2018.2 (2018).

32 Grillner, S. Biological pattern generation: the cellular and computational logic of networks in motion. Neuron 52, 751–766, doi:10.1016/j.neuron.2006.11.008 (2006).

33 Huang, T. et al. Identifying the pathways required for coping behaviours associated with sustained pain. Nature 565, 86–90, doi:10.1038/s41586-018-0793-8 (2019).

34 Powell, R. et al. Inhibiting endocytosis in CGRP(+) nociceptors attenuates inflammatory pain-like behavior. Nat Commun 12, 5812, doi:10.1038/s41467-021-26100-6 (2021).

35 Cameron, D. et al. The organisation of spinoparabrachial neurons in the mouse. Pain 156, 2061–2071, doi:10.1097/j.pain.0000000000000270 (2015).

36 Todd, A. J. Identifying functional populations among the interneurons in laminae I-III of the spinal dorsal horn. Mol Pain 13, 1–19, doi:Artn 1744806917693003 10.1177/1744806917693003 (2017).

37 Allard, J. Physiological properties of the lamina I spinoparabrachial neurons in the mouse. J Physiol 597, 2097–2113, doi:10.1113/JP277447 (2019).

38 Calcutt, N. A. Diabetic neuropathy and neuropathic pain: a (con)fusion of pathogenic mechanisms? Pain 161, S65–S86, doi:10.1097/j.pain.0000000000001922 (2020).

39 Molsted, S. & Eidemak, I. Musculoskeletal pain reported by mobile patients with chronic kidney disease. Clin Kidney J 13, 813–820, doi:10.1093/ckj/sfz196 (2020).

40 Sutherland, J. P. et al. Persistence of a T Cell Infiltrate in Human Ganglia Years After Herpes Zoster and During Post-herpetic Neuralgia. Front Microbiol 10, 2117, doi:10.3389/fmicb.2019.02117 (2019).

41 O’Donnell, M. J. et al. Chronic pain syndromes after ischemic stroke: PRoFESS trial. Stroke 44, 1238–1243, doi:10.1161/STROKEAHA.111.671008 (2013).

42 Xing, G. G., Liu, F. Y., Qu, X. X., Han, J. S. & Wan, Y. Long-term synaptic plasticity in the spinal dorsal horn and its modulation by electroacupuncture in rats with neuropathic pain. Experimental Neurology 208, 323–332, doi:10.1016/j.expneurol.2007.09.004 (2007).

43 Cook, A. J., Woolf, C. J., Wall, P. D. & McMahon, S. B. Dynamic receptive field plasticity in rat spinal cord dorsal horn following C-primary afferent input. Nature 325, 151–153, doi:10.1038/325151a0 (1987).

44 Simone, D. A., Baumann, T. K., Collins, J. G. & LaMotte, R. H. Sensitization of cat dorsal horn neurons to innocuous mechanical stimulation after intradermal injection of capsaicin. Brain Res 486, 185–189, doi:10.1016/0006-8993(89)91293-6 (1989).

45 Thompson, S. W., King, A. E. & Woolf, C. J. Activity-Dependent Changes in Rat Ventral Horn Neurons in vitro; Summation of Prolonged Afferent Evoked Postsynaptic Depolarizations Produce a d-2-Amino-5-Phosphonovaleric Acid Sensitive Windup. Eur J Neurosci 2, 638–649, doi:10.1111/j.1460-9568.1990.tb00453.x (1990).

46 Morisset, V. & Nagy, F. Plateau potential-dependent windup of the response to primary afferent stimuli in rat dorsal horn neurons. Eur J Neurosci 12, 3087–3095, doi:10.1046/j.1460-9568.2000.00188.x (2000).

47 Mitsi, V. et al. RGS9-2--controlled adaptations in the striatum determine the onset of action and efficacy of antidepressants in neuropathic pain states. Proc Natl Acad Sci U S A 112, E5088–5097, doi:10.1073/pnas.1504283112 (2015).

48 Shields, S. D., Eckert, W. A., 3rd & Basbaum, A. I. Spared nerve injury model of neuropathic pain in the mouse: a behavioral and anatomic analysis. J Pain 4, 465–470, doi:10.1067/s1526-5900(03)00781-8 (2003).

49 Decosterd, I. & Woolf, C. J. Spared nerve injury: an animal model of persistent peripheral neuropathic pain. Pain 87, 149–158, doi:10.1016/S0304-3959(00)00276-1 (2000).

50 Norman, G. J. et al. Stress and IL-1beta contribute to the development of depressive-like behavior following peripheral nerve injury. Mol Psychiatry 15, 404–414, doi:10.1038/mp.2009.91 (2010).

